# Characterizing allele-by-environment interactions using maize introgression lines

**DOI:** 10.1101/738070

**Authors:** Zhi Li, Sara B. Tirado, Dnyaneshwar C. Kadam, Lisa Coffey, Nathan D. Miller, Edgar P. Spalding, Aaron J. Lorenz, Natalia de Leon, Shawn M. Kaeppler, Patrick S. Schnable, Nathan M. Springer, Candice N. Hirsch

## Abstract

Relatively small genomic introgressions containing quantitative trait loci can have significant impacts on the phenotype of an individual plant. However, the magnitude of phenotypic effects for the same introgression can vary quite substantially in different environments due to allele-by-environment interactions. To study potential patterns of allele-by-environment interactions, fifteen near-isogenic lines (NILs) with >90% B73 genetic background and multiple Mo17 introgressions were grown in 16 different environments. These environments included five geographical locations with multiple planting dates and multiple planting densities. The phenotypic impact of the introgressions was evaluated for up to 26 traits that span different growth stages in each environment to assess allele-by-environment interactions. Results from this study showed that small portions of the genome can drive significant genotype-by-environment interaction across a wide range of vegetative and reproductive traits, and the magnitude of the allele-by-environment interaction varies across traits. Some introgressed segments were more prone to genotype-by-environment interaction than others when evaluating the interaction on a whole plant basis throughout developmental time, indicating variation in phenotypic plasticity throughout the genome. Understanding the profile of allele-by-environment interaction is useful in considerations of how small introgressions of QTL or transgene containing regions might be expected to impact traits in diverse environments.

**Key Message:** Significant allele-by-environment interactions are observed for traits throughout development from small introgressed segments of the genome.

## Introduction

Many important agronomic traits (i.e. grain yield, flowering time, and plant height) are influenced by a large number of quantitative trait loci spread across the genome. These loci have varying effects on phenotypic variation and the effects can be variable across environments. The integration of all allele-by-environment interactions can lead to a complex genotype-by-environment interaction (G×E) effect (Malosetti et al. 2016). Genotype-by-environment interaction is a major challenge for crop breeding as relative performances, and even rankings, of cultivars grown in different environments are not consistent. Sometimes the varieties that have the highest performance in one environment can have poor performance in other environments. Breeders and growers are often faced with choices between yield stability (consistent moderate performance) and yield potential. Increasing our understanding of the profiles of allele-by-environment interactions can provide an improved framework to document and control for these effects in increasingly unpredictable environments (Allard and Bradshaw 1964; El-Soda et al. 2014; Zobel and Talbert 1984).

Although G×E has been well-documented and studied in plant and animal species, it remains a significant challenge to breeding programs (Crossa 2012; de Leon et al. 2016; Xu 2016; Li et al. 2018). Obtaining a rigorous assessment of G×E effects requires phenotypic evaluation in a wide range of environments and breeder choices about the level of trait instability across environments that will be tolerated. Many G×E studies are focused on traits with important economic value, such as yield and quality, which are usually the end products of plants (Ndhlela et al. 2014; Mohammadi and Amri 2016; Balakrishnan et al. 2016). There have been fewer detailed evaluations of G×E interactions for vegetative traits and yield component traits such as ear morphology. Technological advances in high-throughput automated phenotyping have increased opportunities to collect quantitative data on numerous phenotypic traits at multiple developmental stages in a cost-effective and time saving way (El-Soda et al. 2014; Miller et al. 2017).

Studies on G×E frequently utilize contrasting conditions within controlled environments (i.e. managed stress trials) that include extreme conditions like drought/water stress (Chapman et al. 1997; Ribaut et al. 1997; Ribaut and Hoisington 1998), heat/cold stress (Mohammadi et al. 2016; Thiry et al. 2016), or nutritional stress (Bänziger et al. 1997). These extreme conditions or drastic environmental change often result in significant G×E (Bebber et al. 2013; Trenberth et al. 2013; Xu 2016), though it is not necessarily representative of the normal state of the agriculture systems that often are subject to more subtle environmental variations through the use of intensive management practices. G×E interaction studies that utilize natural environments provide a practical understanding of G×E to breeding programs (de Leon et al. 2016; Elias et al. 2016; Malosetti et al. 2016). Data generated by multi-environment trials, which are trials or experiments carried out in multiple environments or contexts with multiple replications, can be quite useful in documenting the range of G×E effects that can be observed (Yan et al. 2001; Fan et al. 2007; Crossa 2012; Malosetti et al. 2013; Nuvunga et al. 2015; Lado et al. 2016; Mohammadi and Amri 2016).

Genotype-by-environment interaction studies that utilize diverse hybrid or inbred lines are limited in that they are unable to identify genomic regions responsible for varying responses to the environment. These studies also provide minimal insight into potential environmental interactions when transgenes or quantitative trait loci are introgressed into a genome. However, potential for extreme interaction of single loci and environment have been reported (Lukens and Doebley 1999; Guo et al. 2014). In this study we evaluated near isogenic lines (NILs) to investigate allele-by-environment interaction. Our multi-environment trials were conducted under normal growing conditions, which provide valuable information for practical breeding programs. Understanding the profile of allele-by-environment interaction will provide a useful framework for interpreting more complex genome-wide G×E. This information is also useful in considerations of how transgenes for quantitative traits or QTL introgressions might be expected to interact with the environment and impact traits.

## Materials and Methods

### Germplasm

A set of 15 B73-like NILs was selected from a previously described set of B73 × Mo17 NILs that included 186 B73-like genotypes with Mo17 introgressions and 70 Mo17-like genotypes with B73 introgressions (Eichten et al. 2011). The full NIL population was planted in St Paul, MN in Summer 2009 and Summer 2010 in single rows (3.35 m long and 0.76 m apart) and six traits were measured including plant height (measured from the ground to the top of the tassel after flowering), tassel branch number, 50 kernel weight, total kernel weight, cob diameter, and kernel row number (Table S1). A subset of 15 B73-like NILs was selected to be grown in additional environments. The majority exhibited phenotypic differences relative to the B73 recurrent parent for at least one of the traits. The set of selected NILs included B004, B040, B043, B049, B055, B102, B107, B111, B122, B123, B125, B131, B154, B175, and B189.

### Field Phenotypic Evaluation of Selected Near Isogenic Lines

The 15 selected B73-like NILs, Mo17, and five entries of B73 (21 total entries) were planted in 16 environments in Iowa, Minnesota, and Wisconsin in the summer of 2015. The 16 environments were defined by location (5 separate locations) and management practices within location including planting date (early and late planting) and plant density (high density: 70,000 plants ha^−1^ and low density: 20,000 plants ha^−1^). Arlington, WI and Waseca, MN had high and low planting densities, for a total of four environments. Curtiss, IA, Kelly, IA, and St. Paul, MN had a factorial of high and low planting density and early and late planting at each site, for a total of 12 environments (Table S2). Within each location/management environment there were two replications and plots were arranged in a randomized complete block design. In each location, plots were grown as single rows (3.35 m long and 0.76 m apart).

A total of 26 traits were collected, including 15 vegetative traits and 11 yield-related traits (Table S3). Thirteen vegetative traits were measured on six representative plants per plot. These traits included plant height at 14, 21, 28, 35, 42, 49, 56, 63 and 70 days after planting (DAP), plant height at maturity, leaf number above the ear, and leaf number (including senesced leaves) below the ear. Juvenile leaves were marked to allow leaf number including senesced leaves to be counted using previously described methods (Hirsch et al. 2014). Days to anthesis and days to silk were measured on a per-plot basis. Custom computer algorithms executed on Open Science Grid computational resources (Pordes et al. 2007) in a workflow managed by HTCondor software (Thain et al. 2005) were used to quantify eleven ear and kernel traits from digital images as previously described (Miller et al. 2017). Six representative ears per plot were measured. Ear weight and grain weight was an average of the weight of the uppermost ear on each of six representative plants in the plot and cob weight was measured on individual uppermost ears from the six representative plants in the plot. For all traits for which single plant measurements were taken, the same six representative plants were used for all measurements. See Table S4 for raw phenotypic values.

Two replicates of the full panel of NILs were also grown in a randomized complete block design in St. Paul, MN in summer 2015 using a single planting date and density (Fig. S1).

### Sequencing and SNP identification

Genotyping of NILs was performed using the genotyping-by-sequencing (GBS) method (Elshire et al., 2011). Briefly, five seeds of each NIL were planted in the greenhouse and pooled leaf samples were collected for each NIL after ten days of planting. Pooled leaf tissue was immediately frozen in liquid nitrogen and then lyophilized. Genomic DNA was extracted from the lyophilized leaf samples using the BioSprint 96 DNA Plant Kit. The quality and quantity of extracted DNA was evaluated using a QuantiFluor^®^ dsDNA System (Promega), and by running DNA samples on 1% agarose gel. Library preparation and sequencing of the DNA samples were performed at the Institute for Genomic Diversity (IGD) at Cornell University as described by Elshire et al. (2011). Single nucleotide polymorphisms (SNPs) were called from the raw sequence data using the TASSEL GBS Pipeline version 3.0 (Glaubitz et al., 2014). SNPs were filtered for maximum missing percentage (> 20%) and minimum minor allele frequency (< 5%).

Beagle software package version 4.1 (Browning and Browning 2016) was employed to remove SNPs that were heterozygous in either parent or showing no polymorphism between the two parents (B73 and Mo17). Missing data in each NIL was then imputed based on the parental genotype. SNPs in each line were recorded as A (B73 allele), B (Mo17) and H (heterozygous). Due to the relatively high rate of missing data and the potential for sequencing error of data generated by the GBS method, a modification of a previously described non-overlapping sliding window approach was used to evaluate SNPs collectively rather than individually (Huang et al. 2009). A window size of 50 SNPs was set continuously along each chromosome. In each window, A, B and H were assigned values of 0, 1 and 0.5, respectively, and the average score of the 50 SNPs in the window was calculated. Within each window if the average score was less than or equal to 0.25, all 50 loci in that window were coded as A. If the average score was greater than or equal to 0.75, all 50 loci in that window were coded as B. If the average score was between 0.25 and 0.75 all 50 loci in the window were coded as H (Table S5).

### Statistical Analyses

Phenotypic variance of each trait was partitioned into environments, genotypes, and G×E effects with a linear model *y*_*ijk*_ = *μ* + *f*_*k*_ + *e*_*i*_ + *r*(*e*)_*j*(*i*)_ + (*ge*)_*k*×*i*_ + *ε*_*ijk*_ where *y*_*ijk*_ is the phenotype value of the *k*^th^ genotype in *i*^th^ environment of the *j*^th^ replication; *μ* is the phenotypic mean across environments; *f*_*k*_ is the *k*^th^ genotype effect; *e*_*i*_ is the *i*^th^ environment effect; *r*(*e*)_*j*(*i*)_ is the *j*^th^ replication effect nested within the i^th^ environment; (*ge*)_*k*×*i*_ is the interaction effect of the *k*^th^ genotype by the *i*^th^ environment, and *ε*_*ijk*_ is the residual effect. All variables except μ were considered as random effects. Two-way ANOVA with replication was conducted with the R function “aov”, and variance components were estimated using least squares. Each trait was analyzed using this model with the 15 NILs and B73.

One-way ANOVA was used to test the phenotypic difference between high density and low density environments for each NIL using the model *y*_*ij*_ = *μ* + *f*_*i*_ + *ε*_*ij*_, where *y*_*ij*_ represents the phenotypic value of the *j*^th^ observation (j=1,2,…,n_i_) on the *i*^th^ density (*i*=1,2,…,k levels); *μ* is the grand mean; *f*_*i*_ represents the *i*^th^ treatment effect, and *ε*_*ij*_ represents the random error present in the *j*^th^ observation on the *i*^th^ treatment. The analysis was done for each trait for each NIL.

For each trait of each NIL in each environment, the phenotypic difference between the NIL and B73 (calculated as B73 minus NIL) was regressed against the mean of all NILs in that environment. A linear regression was fit using the “lm” function in R. A polynomial regression with degree of 2 was fit for traits without significant linear regression using the “lm” function and “ploy” command in R.

To test for differences between environments by using information across all of the traits, the Mantel test (Mantel 1967; Diniz-Filho et al. 2013), which tests for significant differences between pairs of matrices, was used. In this case, the matrices consisted of the 26 traits by 17 genotypes (15 NILs, B73 and Mo17), with a single matrix for each of the 16 environments (16 total matrices). Each matrix was converted to a pairwise euclidean distance matrix (standardized by mean and standard deviation) (Deza and Deza 2009) between genotypes using the “daisy” function in the “cluster” package in R. Pairwise tests between the 16 environment distance matrices were then conducted using the “mantel.rtest” function (“ade4” package) in R. Permutation numbers for the Mantel test was set to 999. Standard deviation (SD) of the pairwise distances between the 15 NILs across all the environments was calculated as a metric of stability of relationships between NILs on a whole plant basis.

Additive main effects and multiplicative interaction analysis (AMMI) analysis was performed using the “AMMI” function in the R package “agricolae” (De Mendiburu and Simon 2015).

### Code Availability

All code used for the analyses in this manuscript is available at https://github.com/lixx5447/NIL_project.

## Results

### Genomic distribution of introgressions in the selected NILs

The goal of this study was to determine if introgression segments throughout the genome exhibit variation in G×E sensitivity, and what interaction patterns are observed across environments for different introgression lines. To address these questions, we selected 15 NILs from a larger population of NILs that was previously described (Eichten et al. 2011). During the development of the complete NIL population, no selection was imposed, and the introgressed segments in each line were considered random. The subset of 15 NILs selected for this study exhibited higher or lower trait values relative to B73 in grow-outs of the full NIL panel in 2009 and 2010 (Table S1) and the full NIL panel was grown in summer 2015 at a single location and condition in order to compare the full set of traits used in this study (Fig. S1). The data for the 2009/2010 grow-outs were used to select a subset of the NILs that exhibited variation for a certain trait relative to B73, but did not necessarily contain major-effect QTL for these traits. Across the 12 traits, the selected NILs (which are primarily B73 background with a small number of Mo17 introgressed segments) showed a large phenotypic effect (deviation greater than 3 times the standard error in either direction) for one to nine traits (Table S6). Within each trait, however, the 15 selected NILs spanned the range of variation that was observed in the complete population of B73-like NILs (Fig. S1). As such, the 15 selected NILs and the introgressed segments were considered random.

To characterize the introgression segments, the 15 selected NILs and the two inbred parents were genotyped using the GBS method (Elshire et al. 2011). After quality control and imputation, 122,516 SNPs were retained for further analysis for 14 of the 15 NILs (Table S5, Fig. S2). Due to a high rate of missing GBS data for NIL B040, genotypic data generated by comparative genomic hybridization (CGH) in a previous study (Eichten et al. 2011) was used in place of the GBS data for this line. Within each of the NILs there was between three and 12 introgressions. The sum of lengths of all introgression segments in a particular NIL ranged from 66.11 Mb (B175, 3.10% of genome) to 327.44 Mb (B125, 15.34% of the genome) with an average of 164.61 Mb (7.71% of the genome) in the NIL set. The percentage of heterozygous introgression segments across the genome ranged from 0 to 1.25% (26.7 Mb) with an average of 0.34% (7.18Mb). The number of annotated genes introgressed into each NIL ranged from 1,299 to 6,360 with an average of 3,393 (Table S7).

### Significant genotype, environment, and genotype-by-environment interaction effects are observed across traits within the selected NILs

An analysis of variance was conducted with the 15 NILs and the recurrent parent B73 for each of the 26 traits to initially partition the observed variation by genotype, environment, G×E, and residual. For all measured traits, genotype and environmental sources of variation were significant at P < 0.001 (Table 1). In addition, 15 of the 26 traits exhibit significant G×E effects. As expected, the proportion of the variation explained by genotype, environment, and G×E effects varied substantially among the different traits. The percent variation explained by G×E effect ranged from a mere 2.2% (Plant Height at 28 DAP) up to 35.0% (Kernel Depth) of the trait variation. There were nine traits, all related to ear or kernel morphology, for which G×E effects accounted for at least 20% of the total variation. These observations provide evidence for substantial environmental and G×E effects within the set of selected NILs and the environments used in this study.

**Table 1.**
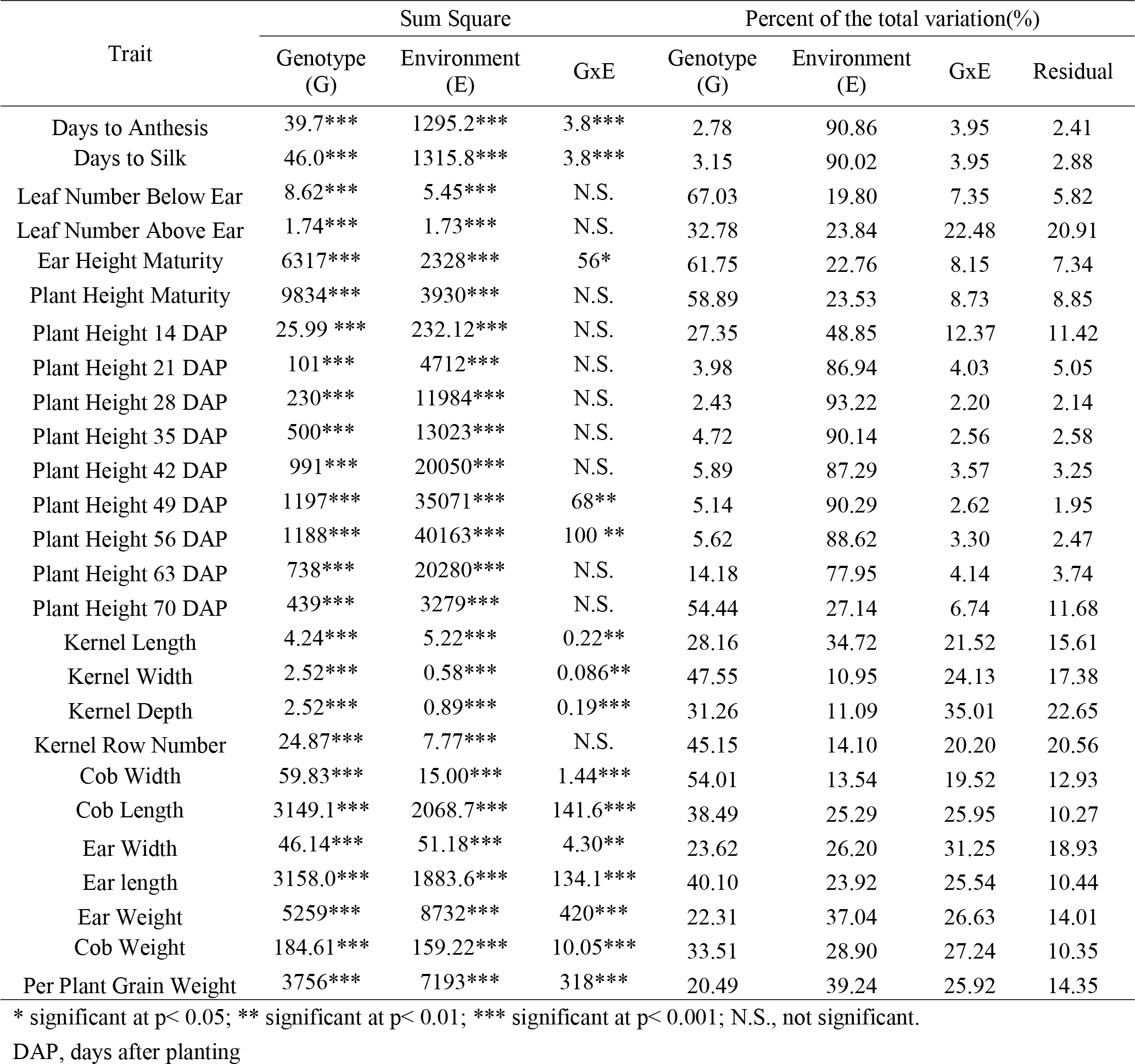
Analysis of variance for 15 near isogenic lines (NILs) and B73 for 26 traits measured in 16 environments.

| Trait | Sum Square |  |  | Percent of the total variation(%) |  |  |  |
| --- | --- | --- | --- | --- | --- | --- | --- |
|  | Genotype (G) | Environment (E) | GxE | Genotype (G) | Environment (E) | GxE | Residual |
| Days to Anthesis | 39.7*** | 1295.2*** | 3.8*** | 2.78 | 90.86 | 3.95 | 2.41 |
| Days to Silk | 46.0*** | 1315.8*** | 3.8*** | 3.15 | 90.02 | 3.95 | 2.88 |
| Leaf Number Below Ear | 8.62*** | 5.45*** | N.S. | 67.03 | 19.80 | 7.35 | 5.82 |
| Leaf Number Above Ear | 1.74*** | 1.73*** | N.S. | 32.78 | 23.84 | 22.48 | 20.91 |
| Ear Height Maturity | 6317*** | 2328*** | 56* | 61.75 | 22.76 | 8.15 | 7.34 |
| Plant Height Maturity | 9834*** | 3930*** | N.S. | 58.89 | 23.53 | 8.73 | 8.85 |
| Plant Height 14 DAP | 25.99 *** | 232.12*** | N.S. | 27.35 | 48.85 | 12.37 | 11.42 |
| Plant Height 21 DAP | 101*** | 4712*** | N.S. | 3.98 | 86.94 | 4.03 | 5.05 |
| Plant Height 28 DAP | 230*** | 11984*** | N.S. | 2.43 | 93.22 | 2.20 | 2.14 |
| Plant Height 35 DAP | 500*** | 13023*** | N.S. | 4.72 | 90.14 | 2.56 | 2.58 |
| Plant Height 42 DAP | 991*** | 20050*** | N.S. | 5.89 | 87.29 | 3.57 | 3.25 |
| Plant Height 49 DAP | 1197*** | 35071*** | 68** | 5.14 | 90.29 | 2.62 | 1.95 |
| Plant Height 56 DAP | 1188*** | 40163*** | 100 ** | 5.62 | 88.62 | 3.30 | 2.47 |
| Plant Height 63 DAP | 738*** | 20280*** | N.S. | 14.18 | 77.95 | 4.14 | 3.74 |
| Plant Height 70 DAP | 439*** | 3279*** | N.S. | 54.44 | 27.14 | 6.74 | 11.68 |
| Kernel Length | 4.24*** | 5.22*** | 0.22** | 28.16 | 34.72 | 21.52 | 15.61 |
| Kernel Width | 2.52*** | 0.58*** | 0.086** | 47.55 | 10.95 | 24.13 | 17.38 |
| Kernel Depth | 2.52*** | 0.89*** | 0.19*** | 31.26 | 11.09 | 35.01 | 22.65 |
| Kernel Row Number | 24.87*** | 7.77*** | N.S. | 45.15 | 14.10 | 20.20 | 20.56 |
| Cob Width | 59.83*** | 15.00*** | 1.44*** | 54.01 | 13.54 | 19.52 | 12.93 |
| Cob Length | 3149.1*** | 2068.7*** | 141.6*** | 38.49 | 25.29 | 25.95 | 10.27 |
| Ear Width | 46.14*** | 51.18*** | 4.30** | 23.62 | 26.20 | 31.25 | 18.93 |
| Ear length | 3158.0*** | 1883.6*** | 134.1*** | 40.10 | 23.92 | 25.54 | 10.44 |
| Ear Weight | 5259*** | 8732*** | 420*** | 22.31 | 37.04 | 26.63 | 14.01 |
| Cob Weight | 184.61*** | 159.22*** | 10.05*** | 33.51 | 28.90 | 27.24 | 10.35 |
| Per Plant Grain Weight | 3756*** | 7193*** | 318*** | 20.49 | 39.24 | 25.92 | 14.35 |

\* significant at $p < 0.05$ ; \*\* significant at $p < 0.01$ ; \*\*\* significant at $p < 0.001$ ; N.S., not significant.
DAP, days after planting

### NILs with different introgression segments respond variably against macro-environmental factors

Given significant G×E effects for most of the traits, we further explored the patterns of interactions that can be observed among environments for the NIL lines. We first looked at patterns within macro-environments defined based on planting density (i.e. high and low planting density). Reaction norm plots were generated for all traits (Fig. S3). We focus here on six representative traits including cob length (CL), cob width (CW), grain weight per plant (GWT), plant height at maturity (PHt), kernel length (KL), and kernel width (KW) (Fig. 1A). For some traits such as CL, CW and KW, the phenotype of all of the NILs was biased toward the recurrent parent B73 and was significantly different from the donor parent Mo17. Other traits such as GWT, KL and PHt exhibited broader ranges of phenotypic variation that extend beyond the parental range. For PHt, planting density had a similar magnitude of positive effect on almost all the NILs, indicating the absence of a genotype-by-density effect. By contrast, GWT exhibited a variety of effects among the NILs with significant negative effects for six NILs and a positive effect for one NIL. The remaining traits (KL, CL, CW and KW) tended to exhibit negative effects relative to plant density although the effect magnitude varied substantially among the different NILs (Fig. 1A). The total number of NILs with significant variation due to planting density was assessed for all traits (Fig. 1B). A wide range was observed across the traits with some traits showing no NILs with significant differences across the density macro environments (i.e. days to anthesis and days to silking), while for others nearly every NIL had an effect and the effect was in the same direction (i.e. plant height and ear height). Interestingly, a number of traits were observed to have NILs with significant differences in both directions (i.e. ear width, ear weight, and grain weight).

**Fig. 1.**
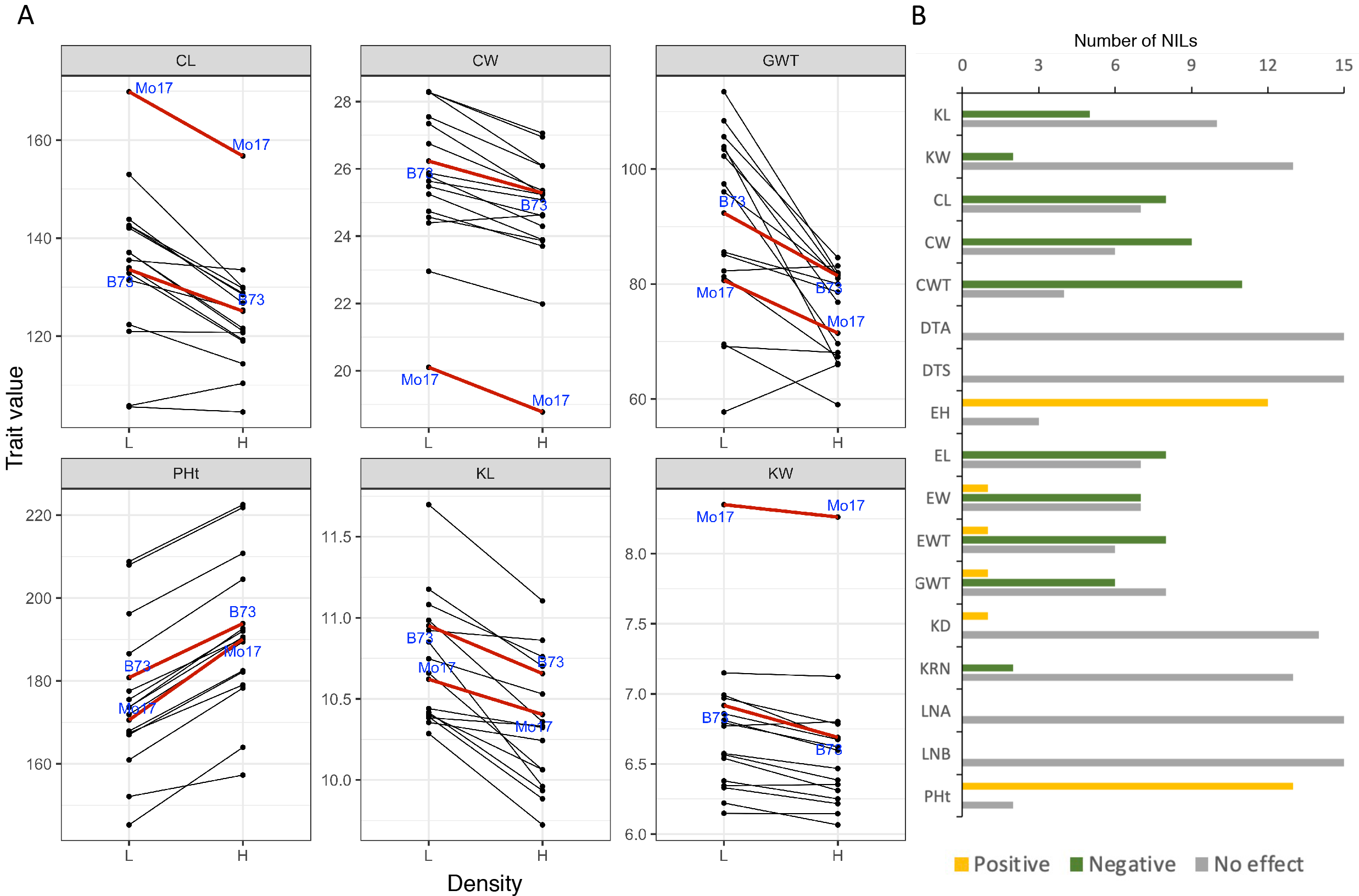
Reaction norm plots of the selected near isogenic lines (NILs) under high and low planting densities. A. Example reaction norms of the NILs under high and low densities for various traits. B. Significance of effect of density on each of the 26 measured traits. Positive and negative effects were defined by a significant one-way ANVOA between the two densities. Positive effect indicates a higher phenotypic value in high density environment. CL, cob length; CW, cob width; GWT, grain weight per plant; PHt, plant height at maturity; KL, kernel length; KW, kernel width; CWT, cob weight; DTA, days to anthesis; DTS, days to silk; EH, ear height maturity; EL, ear length; EW, ear width; EWT, ear weight; KD, kernel depth; KRN, kernel row number; LNA, leaf number above ear; LNB, leaf number below ear.

### NIL introgressions have variable sensitivity to micro-environmental differences and their ability to respond to environmental quality

We next sought to explore the different potential patterns of trait values that were observed among the NILs across individual environments rather than macro-environments. For this analysis the NILs were compared to the trait value of the recurrent parent, B73. The NILs varied substantially in terms of the magnitude, and range, of phenotypic differences relative to the recurrent parent (Fig 2; Fig. S4). In some cases we observed consistent effects of the introgressions across all environments. For example, PHt for B154 and CL for B040 showed very little variation across the environments. While others, such as KL for B043 and GWT in general, showed a wide range of differences across the environments.

**Fig. 2.**
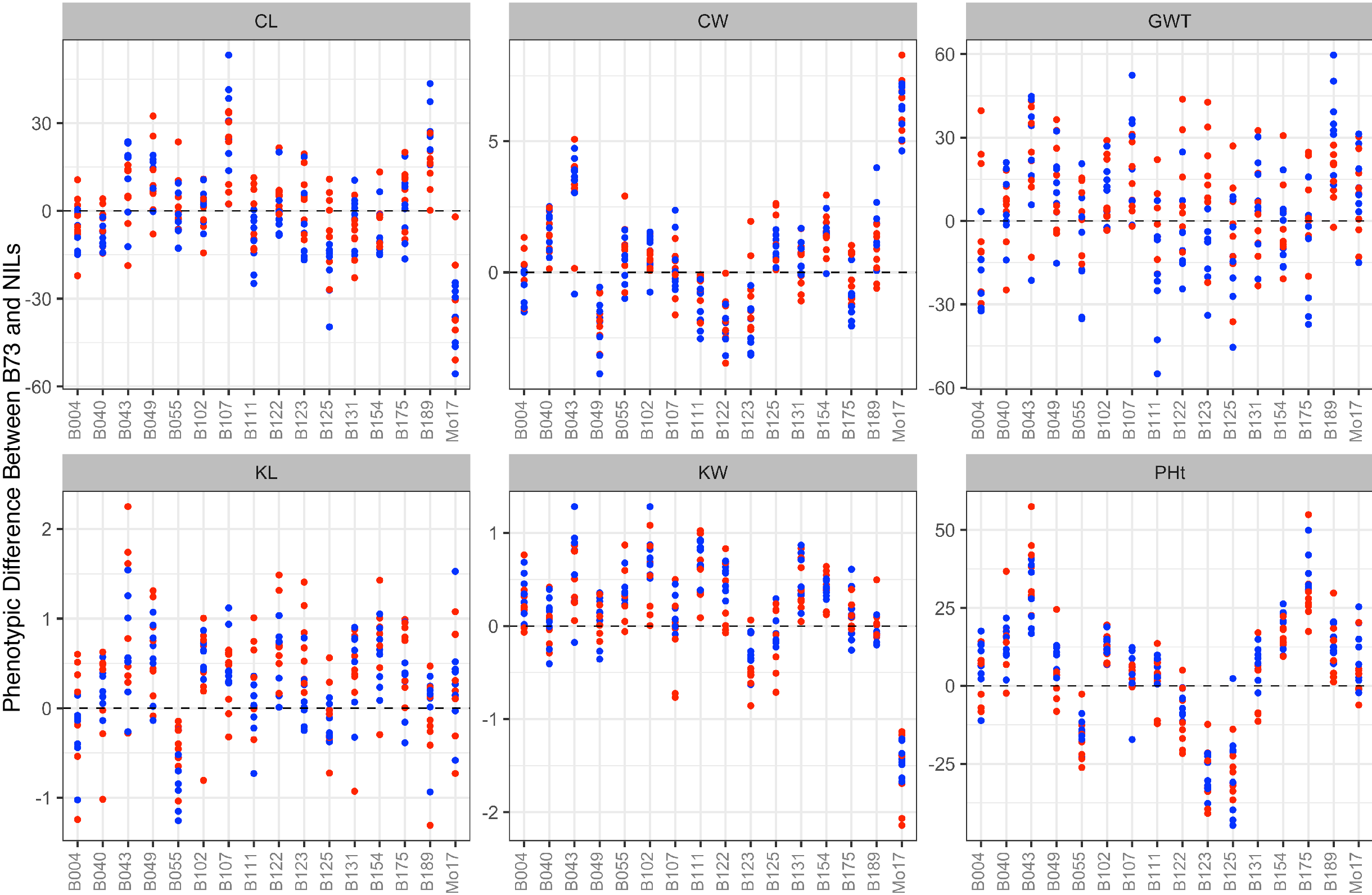
Example distribution pattern of NILs in different environments relative to the recurrent parent. Red and blue dots indicate high and low planting densities respectively. Difference is calculated as B73 minus NIL. CL, cob length (mm); CW, cob width (mm); GWT, per plant grain weight (g); KL, kernel length (mm); KW, kernel width (mm); PHt, plant height at maturity (cm).

We further evaluated the effect of the introgressed segments across the micro-environments by plotting the difference between the NIL and the recurrent parent versus the environmental mean (Fig. 3). We can predict different patterns that might be observed in this type of plot. For example, for some NILs we expect to see a linear relationship in the plot, which would indicate an introgression that has a phenotypic effect and the effect size is directly related to the environmental quality. Indeed, we frequently observe this pattern as shown in Figure 3A (pattern A, corresponding with CL for NIL B189 and KW for NIL B004), in which the introgression effect was positively correlated with environment quality (P = 0.002 and P = 7.9e−05, respectively). We also observed instances of a significant negative correlation between introgression effect and environment quality (pattern B, corresponding with KL for NIL B043 and B111, P = 0.004 and P = 0.009 respectively). Non-linear relationships (P > 0.05 when a linear regression was fitted) were also observed, such as a parabolic-like curve (pattern C, corresponding with PHt for NIL B004 and seed length for NIL B131). However, when a polynomial regression model (degree = 2) was used to fit the data, no significant relationship was detected (P > 0.05), indicating the existence of some other type of non-linear relationship (Fig 3C). Finally, we found some ingressions showed a relatively constant genotype effect regardless of the increase of environmental quality (pattern D, corresponding with CW for NIL B043 and PHt for NIL B175).

**Fig. 3.**
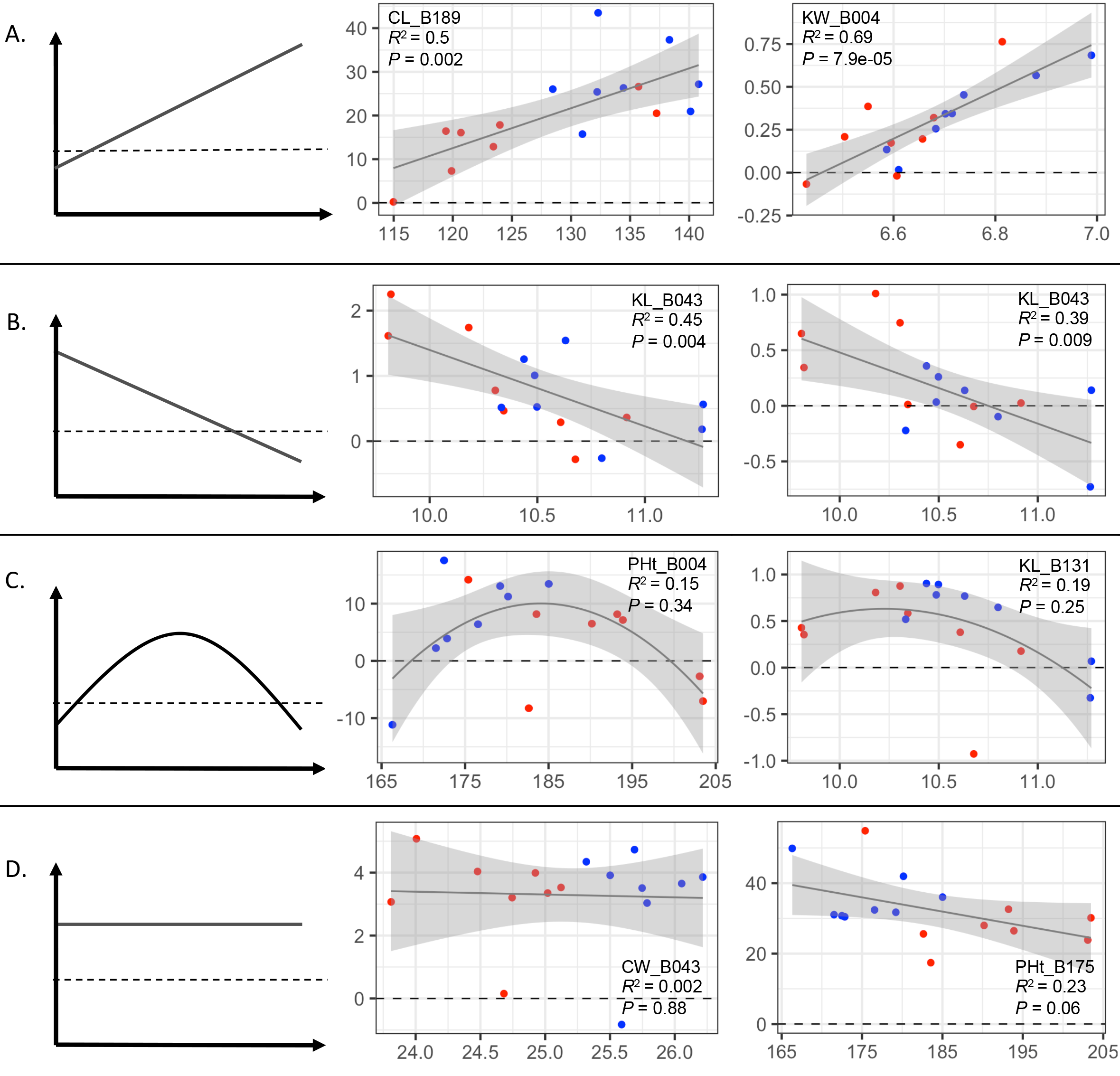
Example phenotypic patterns that are observed for NILs across 16 environments. Data points are arranged on the x-axis according to the mean of all NILs within the environment and the y-axis is the phenotypic difference between the NILs and B73 calculated as B73 minus NIL. Linear regression was fitted for data in panel A, B and D; polynomial regression (degree equals 2) was fitted for data in panel C. The black line or curve in each plot and the nearby gray shaded area indicate the fitted line and its corresponding standard error. The corresponding trait and NIL name as well as R-square and p value for each model are shown in the embedded text. Red and blue dots indicate high and low planting density. CL, cob length; KW, kernel width; KL, kernel length; PHt, plant height at maturity; CW, cob width.

### Environments do not group by geography or planting date or density when environmental similarity is estimated by whole plant performance

The average performance of all genotypes in an environment can provide an estimate of the biological quality or the physical properties of the environment (Finlay and Wilkinson 1963; Malosetti et al. 2013). In the previous analyses the quality of each environment was defined based on the average performance of all individuals in that environment for a single trait. In this study, we collected phenotypic data on traits that span different growth stages, which provided an opportunity to evaluate the relationship between different environments from the perspective of whole plant performance. To do this, a Euclidean distance was calculated between each pair of NILs from all available traits in each environment, which can also be referred to as biological distances (or “biodistances”) (Pilloud and Hefner 2016). The resulting dissimilarity matrix formed by the biodistance between each pair of NILs in each environment was used to define the biological property of that environment. To test the similarity between environments we conducted a Mantel test between each pair of environment dissimilarity matrices. The Mantel test is a commonly used method to evaluate the relationship between geographic distance and genetic divergence (Mantel 1967; Diniz-Filho et al. 2013). This test can assess whether two dissimilarity matrices are correlated with each other. We observed significant correlations between most of the environments based on this test (Fig. 4). However, the magnitude of correlation varied among different pairs of environments. Interestingly, there was limited evidence for consistent clustering of environments by any single factor for which environments could potentially group such as geographic location, planting density, planting date (Fig. 4).

**Fig. 4.**
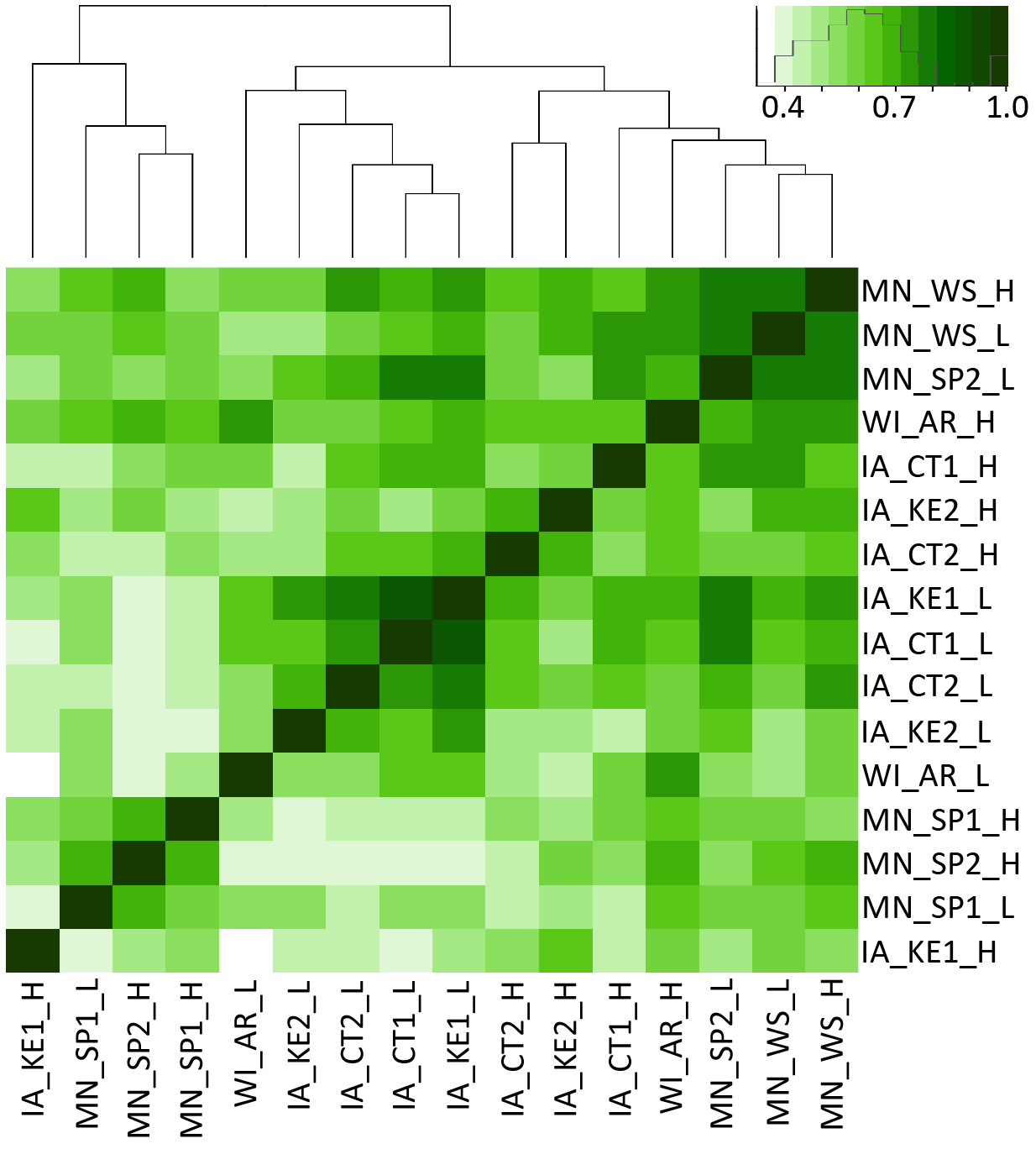
Similarity between environments as determined by similarity of performance of 15 NILs using 26 phenotypic traits. Euclidean distances were calculated between each pair of NILs within each environment based on the phenotypic values of 26 traits. The Mantel test was then run on each pair of matrices across the 16 environments. The heatmap shows the correlation between each pair of environments defined by the results from pairwise Mantel tests. All correlations were significant at p=0.05.

### Combined analysis of all traits suggests different introgressions exhibit different degrees of genotype-by-environment interaction effects

We were also interested to test if there were differences in stability of genotypes with introgressions when evaluating performance on a whole plant basis. To do this we used the biodistances calculated above for each pair of NILs in each of the 16 environments. If there is no allele-by-environment effect from the introgressed segments, the biodistance of a given pair of NILs should be consistent across different environments. We calculated the standard deviation (SD) of the biodistance between each pair of NILs across all 16 environments as an indicator of the biodistance variation. A wide range of biodistances were observed for most of the pairs of NILs (Fig. 5), which indicates that small introgressions show variable stability on a whole plant basis across diverse environments. We found biodistance variation for some NILs were consistently high or low between most of the other NILs. For example, the average SD of the biodistance for NILs B004, B107 and B043 across all the other NILs were 1.78, 1.75 and 1.70 respectively, while for NILs B154, B102 and B040, the values were 1.24, 1.29 and 1.28 respectively (Fig. 5). This result indicates the segments introgressed into some NILs are more or less likely to show allele-by-environment interactions.

**Fig. 5.**
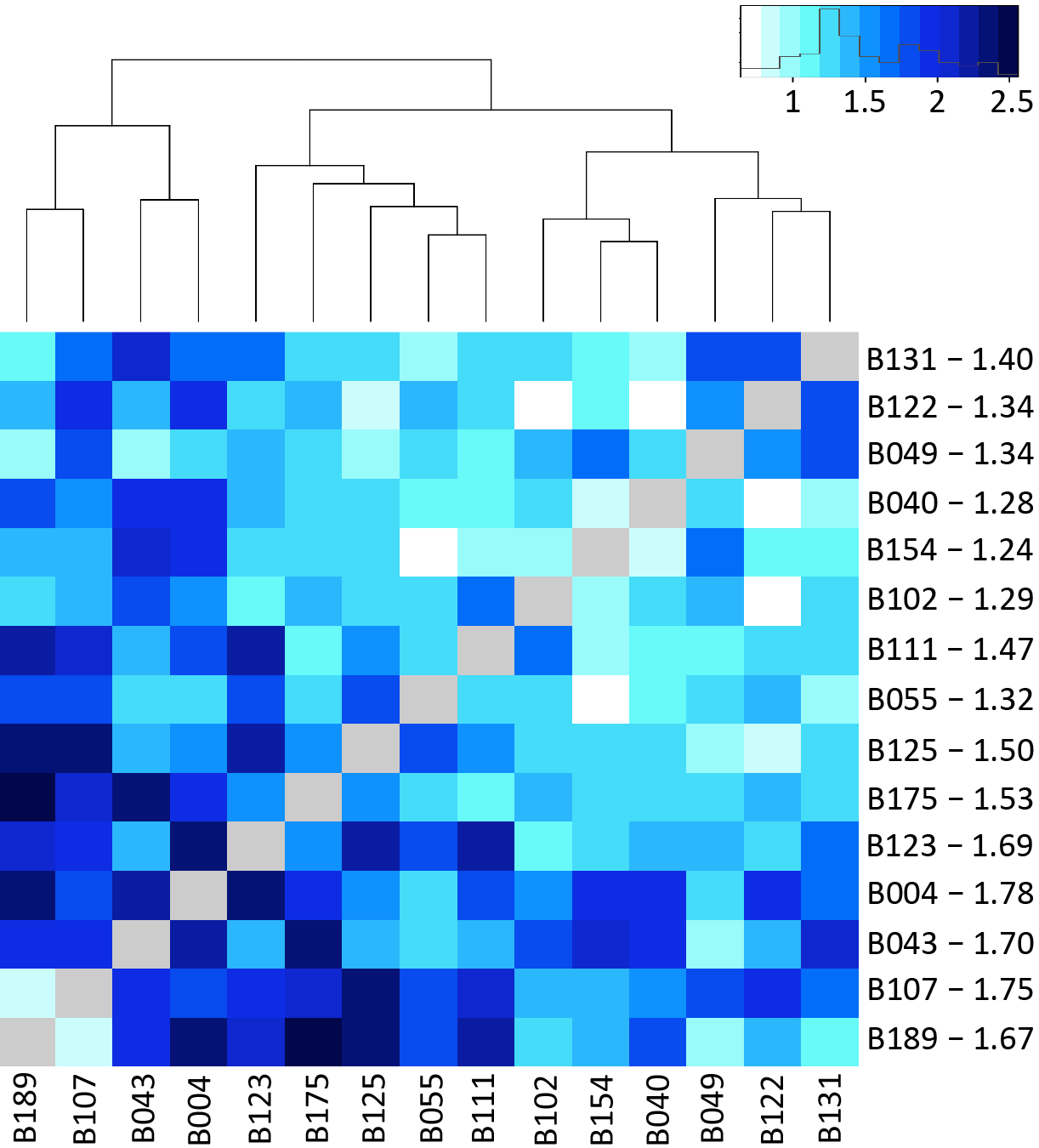
Variation in pairwise relationships between NILs across 16 environments. Standard deviation (SD) of the biodistance between each pair of NILs across all 16 environments was calculated as an indicator of the biodistance variation. Numbers on the right side of the heatmap are the average values of the SD of the biodistance variation for each NIL with all of the other NILs.

We also use the additive main effects and multiplicative interaction analyses (AMMI) model (Gauch 1988; van Eeuwijk 1995) to understand and classify interactions between NILs and environments. When using AMMI models to dissect the G×E for each single trait, we saw different patterns across the distinct traits (Fig. S5). The relative stability of genotype-environment combination varied across different traits. The relationship between environments for single trait analysis can be very different from the results obtained from the Mantel test (Fig. 5), which is on the whole plant basis. For example, B004 is one of the most interactive NILs on the whole plant basis, however, it is very stable in the AMMI results (near the center of the biplot) for five out of the six investigated traits. While for some NILs, such as B043 and B040, which are among the most interactive and stable NILs respectively, the results of AMMI analysis are consistent with that from the analysis done on an entire plant basis (Fig. 5, Fig. S5).

## Discussion

Genotype-by-environment interactions can be highly complex. In this experiment we reduced this complexity by focusing on the genotype, environment, and G×E effects of small introgressed segments of the genome. Introgression of small parts of a genome are common in plant breeding programs to backcross in transgenes or other naturally occurring QTL for traits of importance such as disease resistance. These introgressed segments do not always perform as expected in different backgrounds or environments. Here, we tested to what extent small introgressed segments impact phenotypes and drive G×E to provide a foundation to interpret the results of introgressions of transgenes or QTL. We observed significant impacts of these small introgressions across a wide range of phenotypic traits and on a whole plant basis and found that the effect size and the allele-by-environment interaction of different introgressed segments vary substantially.

The significant effects that we observed were underscored by a number of different patterns of interactions. Introgressed regions that showed a consistent effect across environments, such as cob width for NIL B043 and plant height at maturity for NIL B175, could be defined as constitutive, and are a main target for breeding programs because of their relatively stable effect in all potential growing regions (Fig. 3, pattern D). However, most introgressions (and potential QTL within those introgressions) conferred environment-specific effects on traits, and likely contribute to observed QTL-by-environment interaction effects (Fig. 3, pattern A-C). Those environment-specific QTL can still be very useful for regional breeding if the QTL-by-environment effect can be exploited to achieve a superior effect in adapted environments without a penalty in other environments. Some introgressed regions had opposing phenotypic effects in different environments (such as NIL B175 for ear weight and NIL B049 for kernel row number in Fig. S4). These interactions are relevant to breeders, as they can select the preferred allele in their targeted environment. The QTLs within introgressed regions in these lines likely caused antagonistic pleiotropic effects (El-Soda et al. 2014), but the results could also be due to contrasting effects of different loci within the introgression regions. We also found some introgressed regions that had effects in several environments, but not in others, similar to conditional neutrality (Holland 2007; Zhao and Xu 2012; Tuberosa 2012) (Fig. S4). It is important to remember, that the NILs used in this study contained an average of ~2,800 introgressed genes, and the phenotypic difference between the recurrent parent B73 and the NILs in this study was a combined effect of all the potential QTL/genes in the introgressed regions. However, considering the diverse G×E effects brought by relatively small number/size of introgression segments, it is an important consideration in thinking about how transgenes for quantitative traits or quantitative trait loci introgressions might be expected to impact traits in a wide range of environments.

Previous reports have shown that genotypes can have variable response and tolerance to planting densities (Mansfield and Mumm 2014). In this experiment we included 16 environments that could be divided into macro environments based on planting density. We observed different patterns of reaction norms across both genotypes and traits including grain weight (Fig. 1). Gonzalo et al (2010) detected a QTL for response to planting density for maize grain yield on chromosome 4 in a RIL population derived from the cross of B73 × Mo17. They observed a significant effect difference when replacing the B73 allele with the Mo17 allele at this locus under different planting densities for grain yield. The allele from Mo17 had significantly higher effect under low planting density compared with high density. In our NIL set, there were four NILs (B004, B040, B055, B154) that had introgression segments from Mo17 in this QTL region, and three of them (B004, B055, B154) had significantly higher grain yield in low density environments (Fig. 1), which is consistent with the results of Gonzalo et al.

Maize grain yield is a complex quantitative trait controlled by many small-effect genetic factors (Holland 2007). The continual increase of planting density has been an important factor for genetic yield gain of maize hybrids over time (Duvick 2005a; Messina et al. 2009; Edwards 2016). However, increasing planting density induces stresses upon plants as they compete for resources such as sunlight, water and nutrients in the soil (Mansfield and Mumm 2014). Previous studies have shown that older hybrids have as much yield potential as newer elite hybrids when grown under stress free conditions (Duvick 2005b), however, little is known about the density tolerance of maize inbred lines. In this study, we found that almost all of the NILs have higher grain yield production under lower stress environments (low density) except B189 (Fig. 1A). NIL B189 has two heterozygous introgression segments on chromosome 2 and 5 with a total length of ~24.7Mb and four homozygous introgression segments from Mo17 on chromosome 1, 5, 6 and 10 with a total length of 178.5Mb (Fig. S2 and Table S7). It is not clear whether this increasing density tolerance came from epistatic effects between the homozygous introgression segments from Mo17 and other genomic regions of B73, or the heterotic effect of the heterozygous segments. Further studies on these introgression segments in NIL B189 could provide interesting and useful information about the genetic control of maize density tolerance mechanism.

G×E is complex and there have been many different approaches to both understand and exploit and/or manage this interaction. However, the majority of these studies have been focused on G×E where the genotype of each individual in the study is largely different. Here we asked, what is the degree and patterns of G×E that can be expected with small introgressed segments of a genome? This study provides an important framework for developing breeding strategies when backcrossing in QTL and/or transgenes and the types and patterns of effects that are possible.

## Author contribution statement

CNH, NMS, PSS, SMK, and NdL conceived and designed the experiment. ZL, SBT, LC, NDM, and EPS conducted phenotypic data collection and analysis. ZL, DCK, and AJL performed genotypic data analysis. ZL, NMS, and CNH wrote the manuscript. All authors read and approved the final manuscript.

## Acknowledgement

We thank Jacob Garfin, Peter Hermanson, James Satterlee, Kimberly McFee, and Brad Keiter for technical assistance. This work was supported by the Minnesota Corn Research and Promotion Council (Project Number 4108-16SP), the Minnesota Agricultural Experiment Station (Project 13-113), the Iowa Corn Growers, and Iowa State University’s Plant Sciences Institute.

## Conflict of Interest

On behalf of all authors, the corresponding author states that there is no conflict of interest.

