## Supplemental Figures 1-5 for "Characterizing allele-by-environment interactions using maize introgression lines"

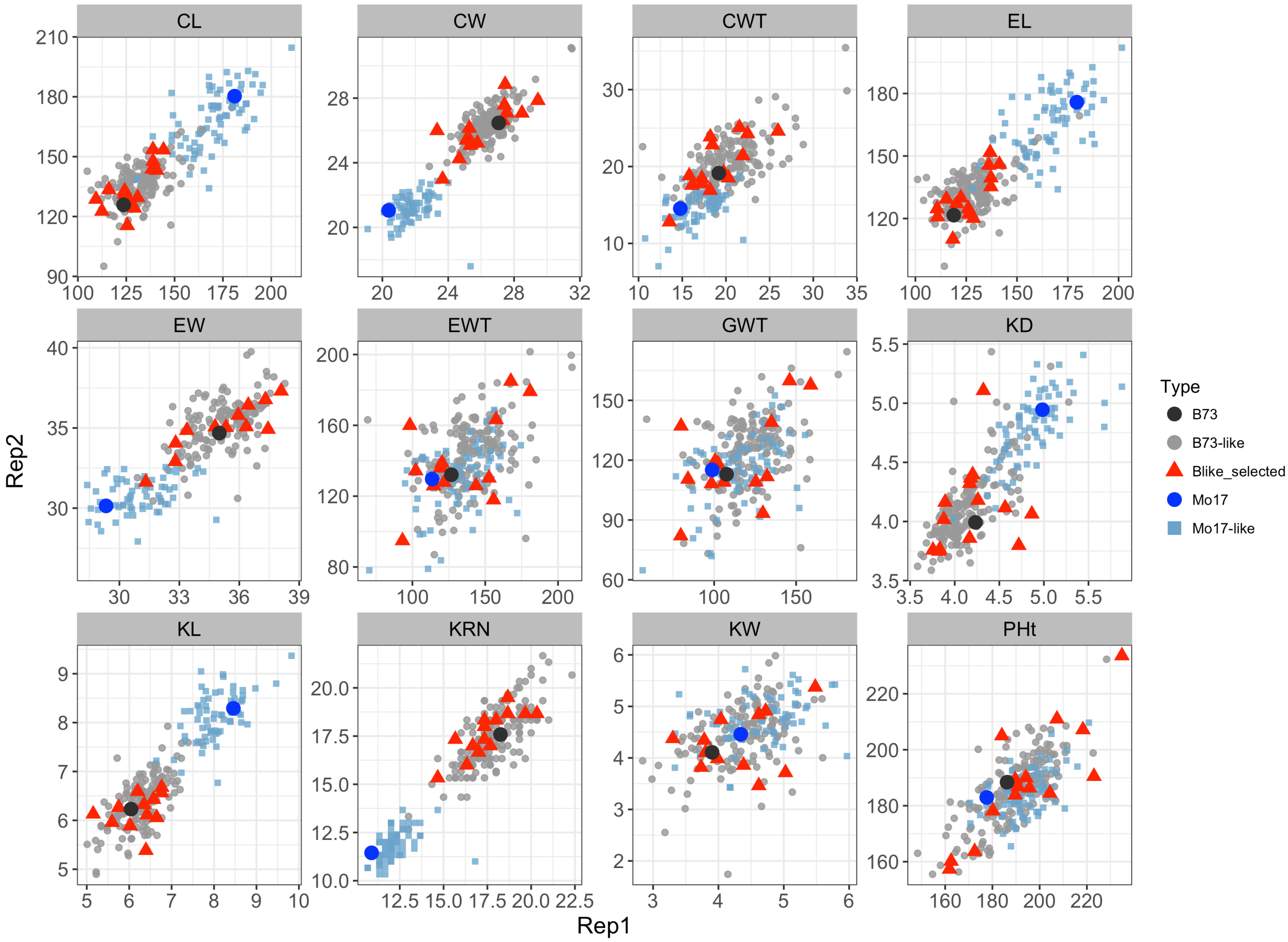


**Figure S1. Phenotypic distribution of the complete B73-Mo17 Near Isogenic Line (NIL) population and selected NILs.** B73-like NILs have B73 as the recurrent parent with Mo17 segements intogressed . Mo17-like NILs have Mo17 as the recurrent parent with B73 segments introgressed. All selected NILs are B73-like NILs. CL, cob length; CW, cob width; CWT, cob weight; EL, ear length; EW, ear width; EWT, ear weight; GWT, per plant grain weight; KD, kernel depth; KL, kernel length; KRN, kernel row number; KW, kernel width; PHt, plant height at maturity.

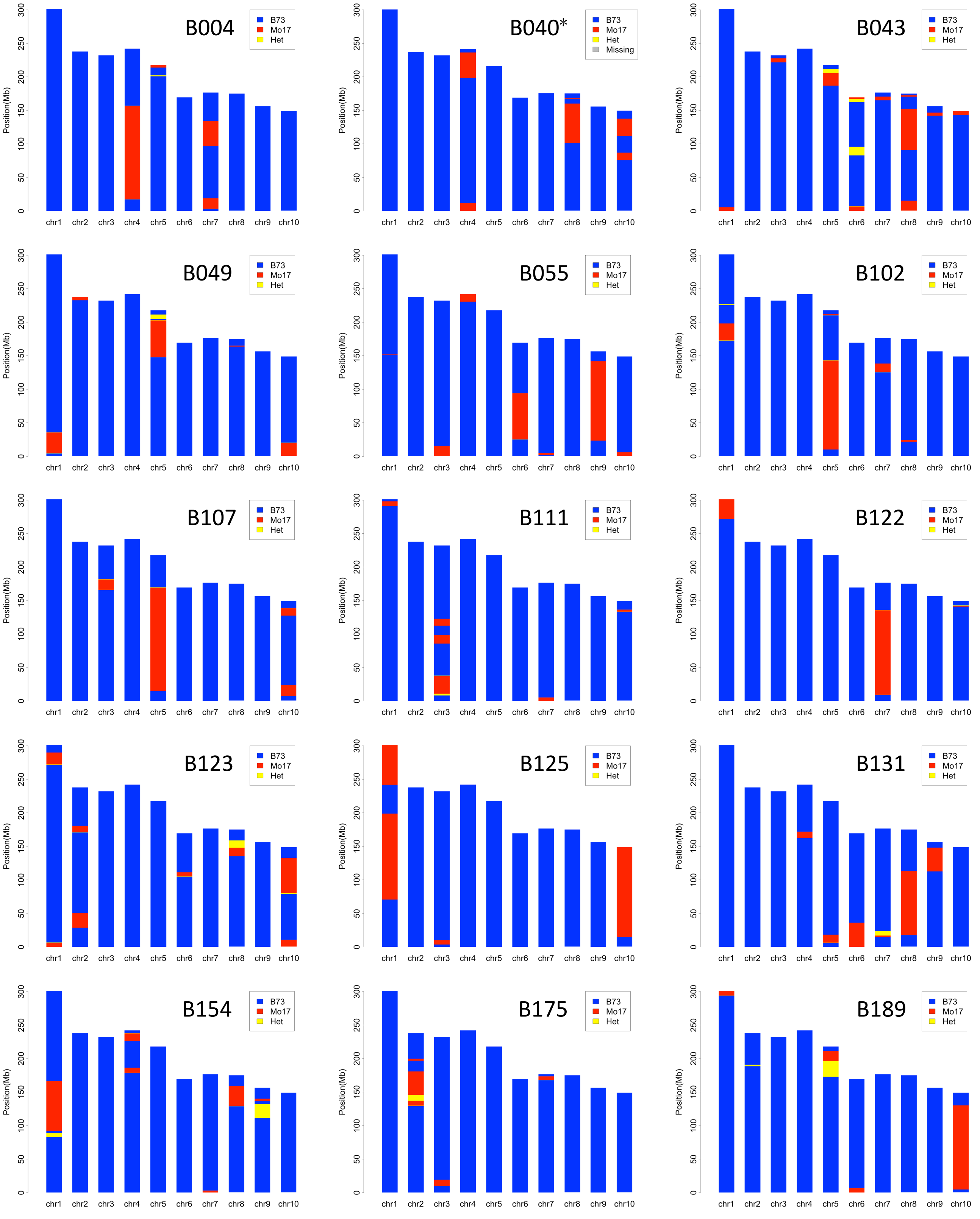


**Figure S2. Distribution of Mo17 introgressions in the selected B73-like near isogenic lines.** Genotyping was performed using genotype-by-sequencing for all lines except B040 which was determined by comparative genome hybridization.

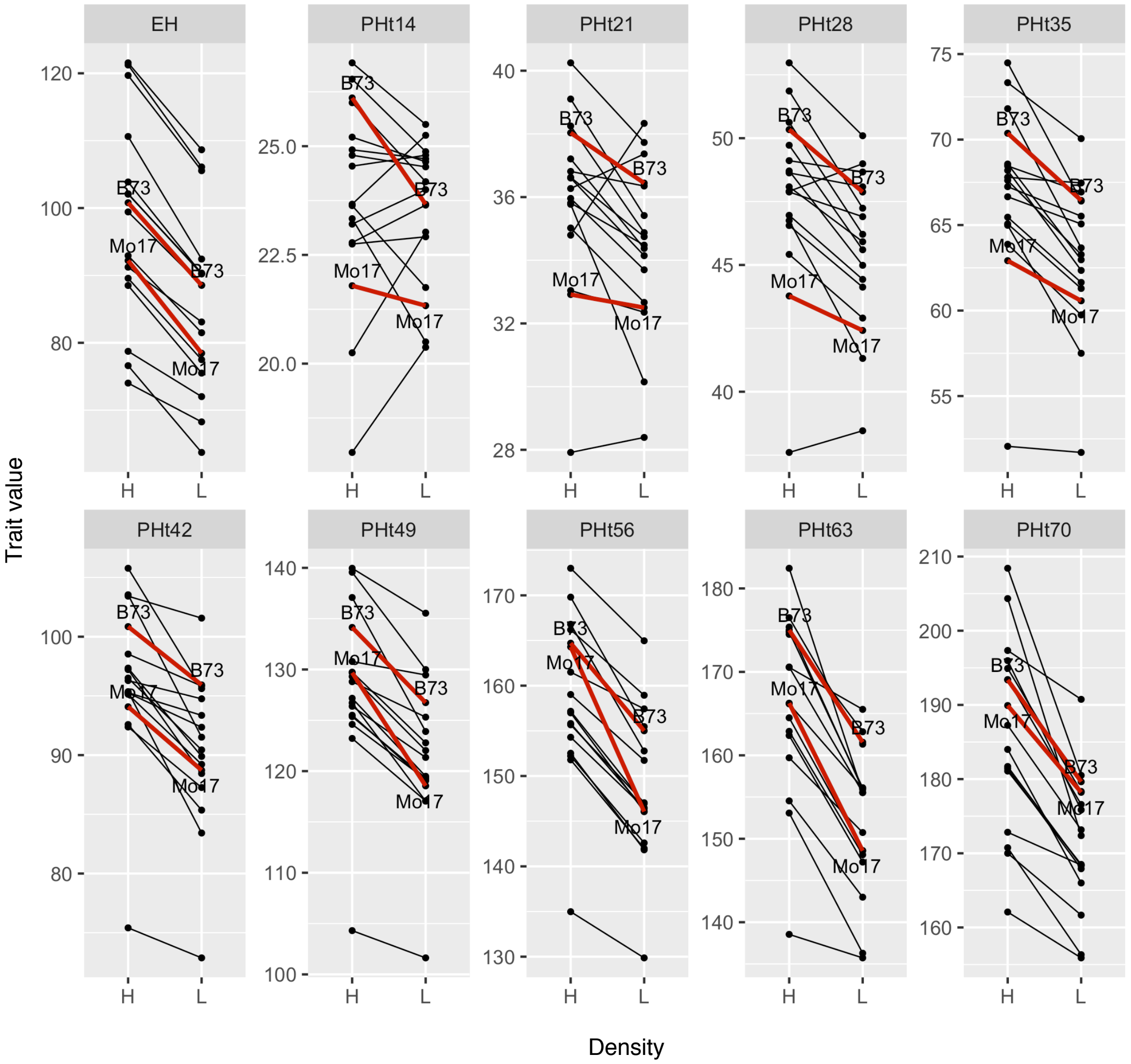


**Figure S3 Part I. Reaction norm plots of the selected NILs under high and low planting densities.** EH, ear height maturity; PHt14, plant height 14 days after planting (DAP); PHt21, plant height 21 DAP; PHt28, plant height 28 days after planting; PHt35, plant height 35 DAP; PHt42, plant height 42 DAP; PHt49, plant height 49 DAP; PHt56, plant height 56 DAP; PHt63, plant height 63 DAP; PHt70, plant height 70 DAP.

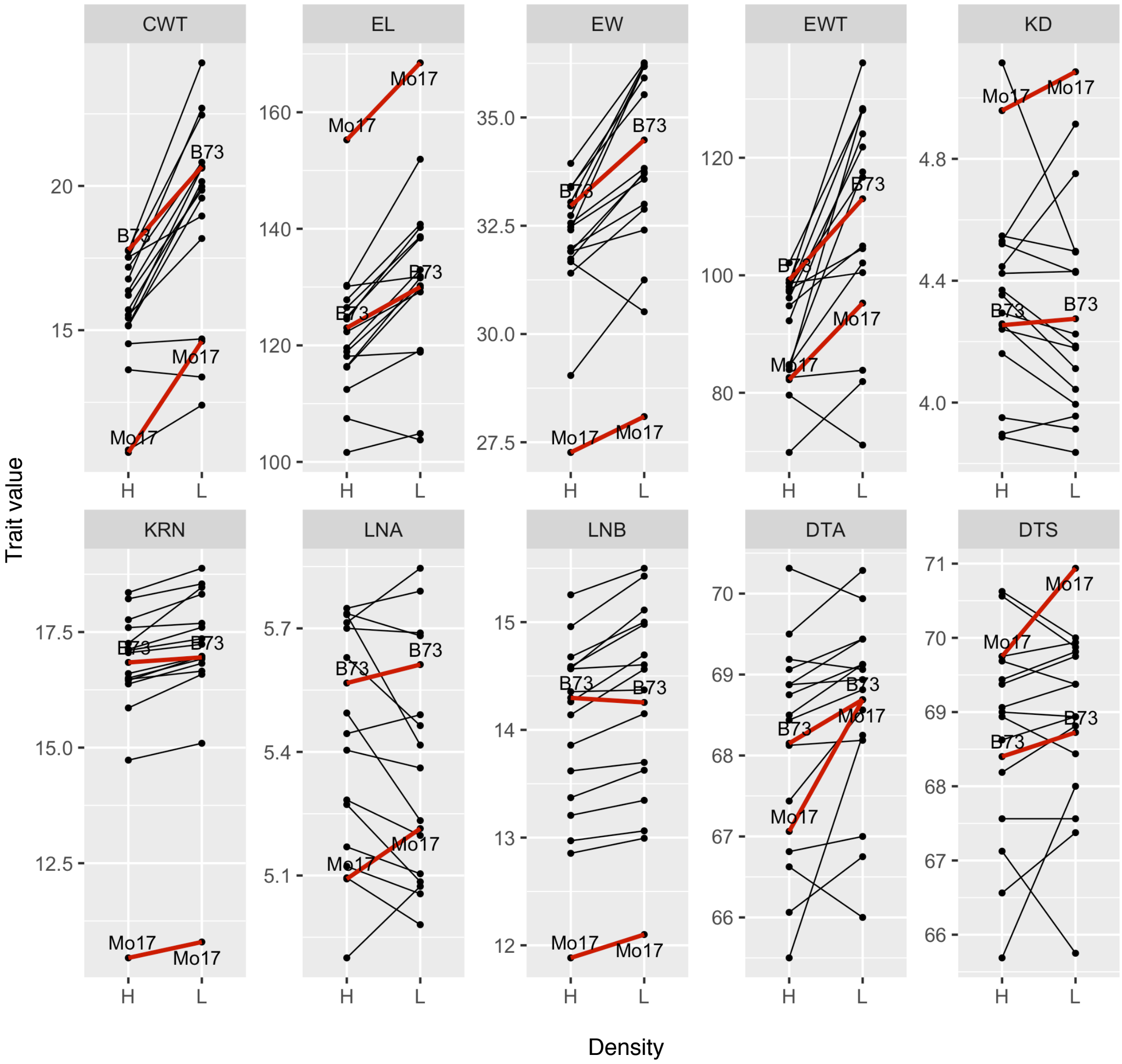


**Figure S3 Part II. Reaction norm plots of the selected NILs under high and low planting densities.** CWT, cob weight; EL, ear length; EW, ear width; EWT, ear weight; KD, kernel depth; KRN, kernel row number; LNA, leaf number above ear; LNB, leaf number below ear; DTA, days to anthesis; DTS, days to silk**.**

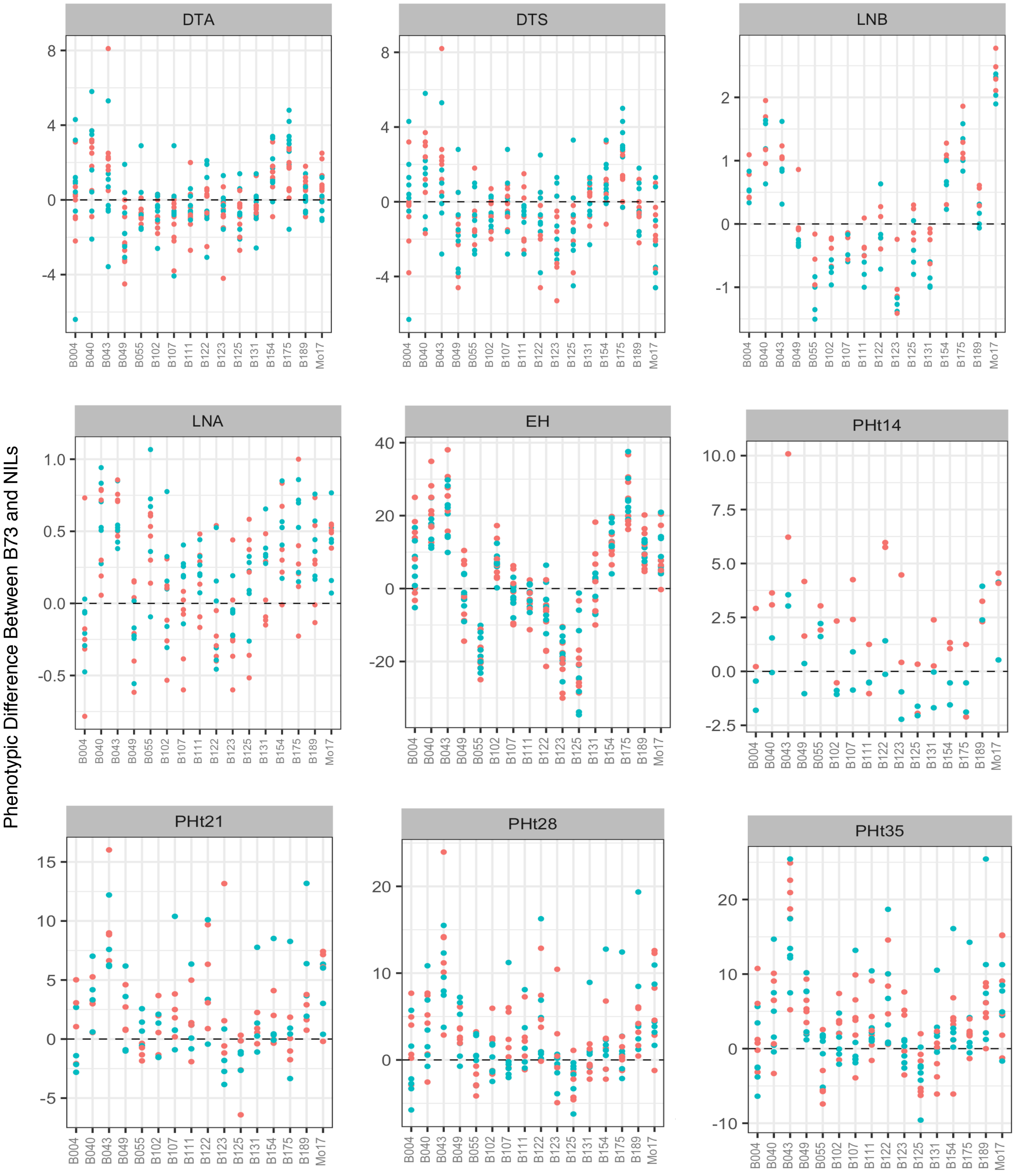


**Figure S4 Part I. Distribution pattern of Near Isogenic Lines (NILs) in different environments relative to the recurrent parent.** Orange and blue dots indicate high and low planting densities respectively. Difference is calculated as B73 minus NIL. DTA, days to anthesis; DTS, days to silk; LNB, leaf number below ear; LNA, leaf number above ear; EH, ear height; PHt14, plant height 14 days after planting (DAP); PHt21, plant height 21 DAP; PHt28, plant height 28 DAP; PHt35, plant height 35 DAP.

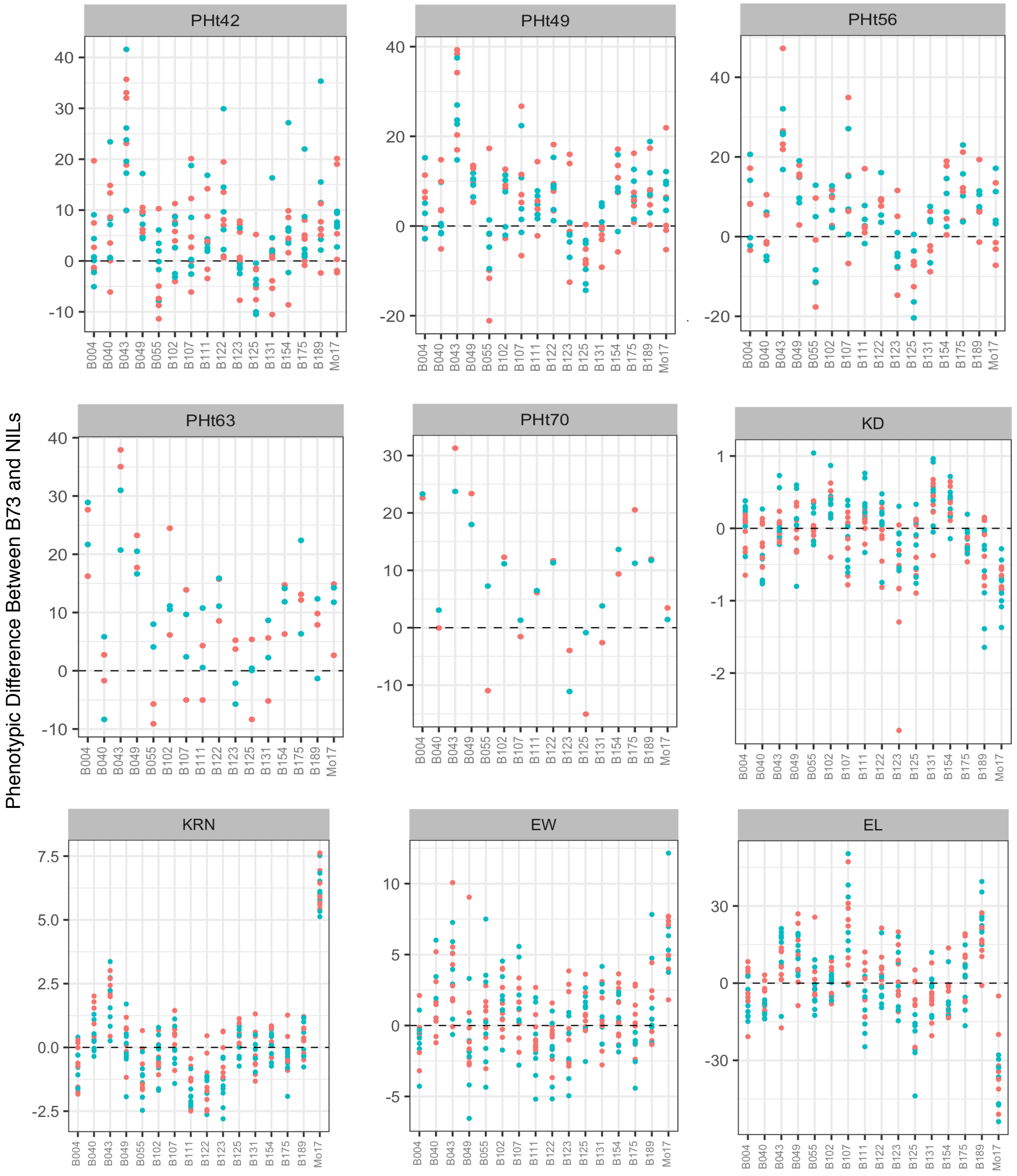


**Figure S4 Part II. Distribution pattern of Near Isogenic Lines (NILs) in different environments relative to the recurrent parent.** Orange and blue dots indicate high and low planting densities respectively. Difference is calculated as B73 minus NIL. PHt42, plant height 42 days after planting (DAP); PHt49, plant height 49 DAP; PHt56, plant height 56 DAP; PHt63, plant height 63 DAP; PHt70, plant height 70 DAP; KD, kernel depth; KRN, kernel row number; EW, ear width; EL, ear length.

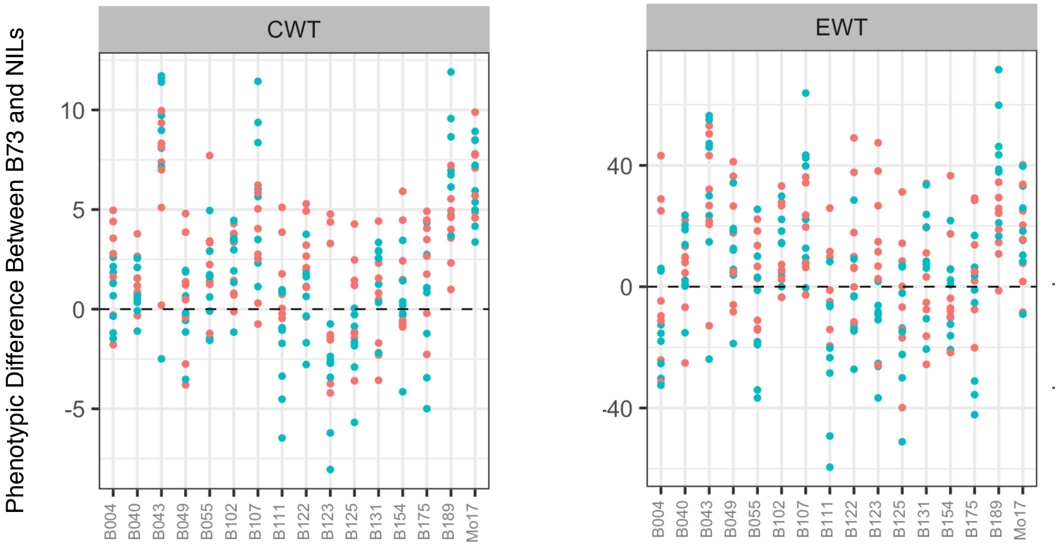


**Figure S4 Part III. Distribution pattern of Near Isogenic Lines (NILs) in different environments relative to the recurrent parent.** Orange and blue dots indicate high and low planting densities respectively. Difference is calculated as B73 minus NIL. EWT, ear weight; CWT, cob weight.

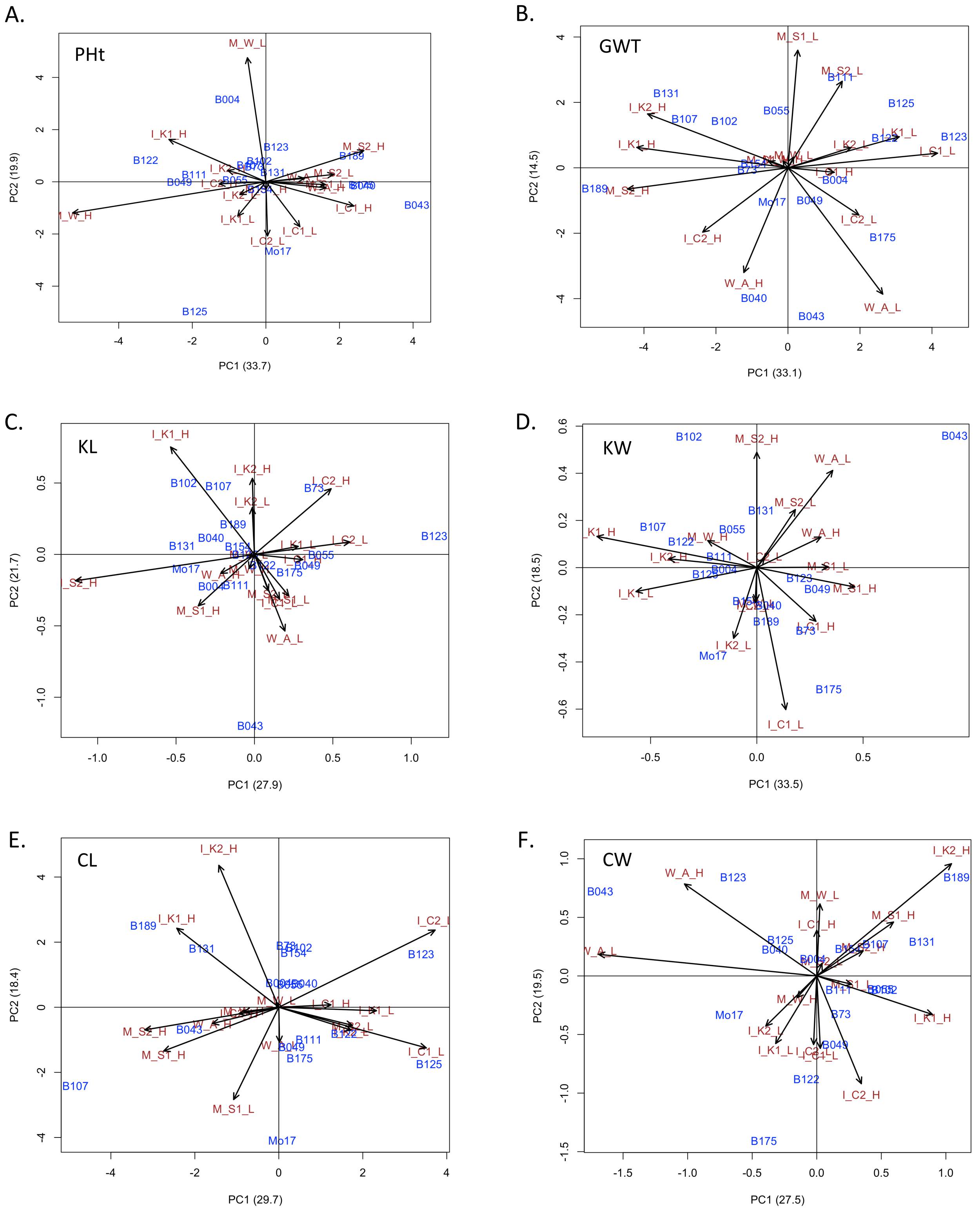


**Figure S5 Additive main effect and multiplicative interaction (AMMI) analysis of GxE.** Traits plotted are A) plant height at maturity, B) grain weight, C) kernel length, D) kernel width, E) cob length, F) cob width. Environments are named as state_location-by-planting_density.
